# piRNA loss unleashes episodic transposition bursts

**DOI:** 10.64898/2026.07.11.737971

**Authors:** Peiwei Chen, Peiyu Xu, Hamna Waseem, John McCormick, Katherine C. Pan, Alexei A. Aravin, Andrew G. Clark, Cédric Feschotte

## Abstract

In Metazoa, transposon expression is suppressed by the piRNA pathway, and disruption of this pathway leads to rampant transposon expression. However, it remains unclear whether increased transposon expression results in actual transposition, and if so, which and how frequently transposons mobilize upon piRNA loss. Here, we developed a framework to track transposon copy accumulation across generations on a single set of nonrecombining haploid genome in the *Drosophila* male germline, with or without the piRNA biogenesis factor HP1D/Rhino. Single-fly Nanopore DNA sequencing revealed that multiple transposon families mobilized after 10-45 generations of piRNA loss. Among them, the most prolific across all replicates was *copia*, producing dozens of new insertions that were distributed across chromosome arms. With genomic DNA collected at every generation, we validated and dated each *copia* insertion, revealing episodic bursts of transposition that deviated strongly from a Poisson process. Using phylogenetic analysis, we further showed that multiple *copia* loci—both autonomous and nonautonomous copies—can mobilize. Interestingly, two additional transposon families, *mdg3* and *invader3*, also mobilized episodically, but with distinct timing and magnitude, highlighting the stochastic nature of transposition bursts. Because we did not observe a single transposition event for these elements in controls, our results provide direct evidence that the piRNA pathway tightly suppresses germline transposition. More importantly, our findings argue that disruption of the piRNA pathway does not simply elevate transposition rates, but it instead unleashes punctuated and stochastic bursts of transposon movement that may radically reshape the timing and magnitude of mutational input during evolution.

**SIGNIFICANCE STATEMENT:** Virtually all animal genomes contain transposons and a small RNA system active in their germline called the piRNA pathway, which is thought to suppress transposition. While it is clear that piRNAs target transposon mRNAs, very few study has explicitly shown that the pathway actually prevents transposition. Rather, the common belief that piRNAs safeguard the germline from transposition comes largely from studies showing that loss of piRNAs leads to increased transposon mRNA expression, but they do not monitor transposon movement. Thus, it remains unclear whether piRNAs broadly control transposition—a central question in the field since the discovery of piRNAs. Here, we follow defined chromosomes through single germline lineages to track transposition and find that piRNA loss unleashes episodic transposition bursts.

## INTRODUCTION

Almost every eukaryotic genome contains potent mutagens: transposons (1). These mobile genetic elements encode or exploit different protein machinery to copy themselves at new genomic loci, generating large-effect mutations and DNA damage. In animals, transposon expression is suppressed by several mechanisms, most notably the germline piRNA pathway (2). Transposon-targeting piRNAs are found in reproductive tissues across diverse metazoans, and disruption of the piRNA pathway often leads to rampant transposon expression (3–5) (Fig. 1A). These observations have led to the prevailing view that the piRNA pathway protects genome integrity by silencing transposons.

**Fig. 1.**
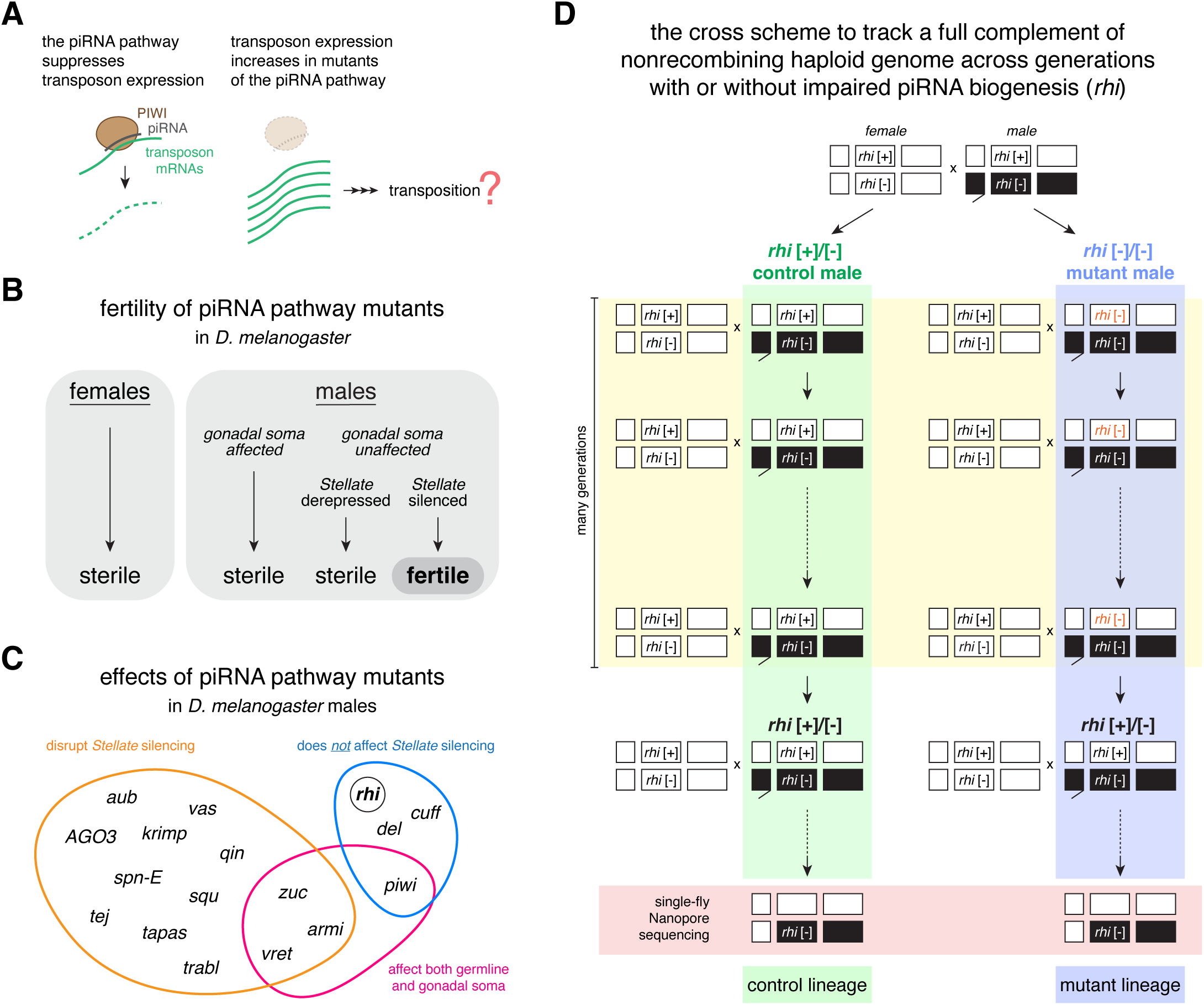
A mutation accumulation strategy for studying transposition with or without Rhi-dependent piRNA biogenesis. (A) A schematic showing the current view of the piRNA pathway function. (B) Summary of the fertility of piRNA pathway mutants in *D. melanogaster*. (C) Summary of the effects of piRNA pathway mutants in *D. melanogaster*, in terms of *Stellate* silencing and cell types affected. (D) The cross scheme to track a full complement of nonrecombining haploid genome across generations with or without *rhi*. Each box represents a chromosome. Filled boxes are focal chromosomes tracked across generations. Because the focal chromosome 2 carried a *rhi* [-] mutant allele, the genotype of each focal male depended on which maternal *rhi* allele it inherited: inheritance of the *rhi* [+] allele produced a heterozygous control male (*rhi* [+]/[-]), whereas inheritance of the *rhi* [-] allele produced a homozygous mutant male (*rhi* [-]/[-]). See also SI Appendix, Fig. S1.

Yet a central assumption underlying this model remains largely untested. The consequences of piRNA pathway disruption have been assessed primarily by measuring levels of transposon RNAs, not transposition itself. As a result, the biological importance of the piRNA pathway is generally inferred from its ability to suppress transposon mRNA transcripts, although the strongest evidence that transposons are deleterious comes from their ability to generate new insertions and mutations. Previous studies have sought to address whether piRNA pathway disruption actually triggers transposition (6–13), using approaches such as transgenic reporters (6) and pooled population sequencing (7–11), typically over short observational windows (7, 8, 12, 13). However, these methods do not directly track new endogenous insertions within defined germline lineages over time, leaving the extent, identity, and dynamics of transposition poorly resolved. Consequently, fundamental questions remain unresolved: to what extent does the piRNA pathway actually prevent transposition? Which transposons mobilize upon piRNA loss? What are the patterns and temporal dynamics of their movement?

## RESULTS

### Identifying an optimal perturbation to study transposition upon piRNA loss

A key challenge in detecting and quantifying transposition following piRNA pathway disruption is that animals lacking an intact piRNA pathway are often infertile (3–5, 14–19). Without offspring, it is difficult to quantitatively assess germline transposition. In the fruit fly *Drosophila melanogaster*, where much of the foundational work on the piRNA pathway has been done (3, 19–21), females with piRNA pathway mutations are typically sterile. In contrast, mutant males are fertile in several cases despite lacking an intact piRNA pathway (Fig. 1B). Motivated by this observation, we sought to identify an optimal perturbation of the piRNA pathway in male flies for studying transposon movement. In *D. melanogaster* gonads, the piRNA pathway functions in both germline cells and gonadal somatic cells, which support germline development (17, 18). Loss of piRNAs in gonadal somatic cells often indirectly disrupts germline development, resulting in infertility. In addition, piRNAs in the male germline silence not only transposons but also the meiotic driver *Stellate* (22, 23), whose derepression can distort inheritance and cause sterility (24–28). Therefore, an ideal perturbation for studies of transposition would disrupt transposon silencing specifically in the germline while preserving *Stellate* silencing as well as male fertility (Fig. 1B).

Examining all known piRNA pathway proteins, about 17 of which have been studied in male flies, we found that only HP1D/Rhino (Rhi), Deadlock (Del), and Cutoff (Cuff) satisfy these criteria (Fig. 1C). These three proteins function together in piRNA biogenesis as an interdependent tripartite Rhi-Del-Cuff (RDC) complex that licenses the transcription of piRNA precursors, and loss of any one component destabilizes the other two (29, 30). Notably, while RDC is required for the expression of most transposon-targeting piRNAs, piRNAs that target *Stellate* are expressed independently of RDC. Consistent with this, *rhi* mutations have been shown to cause severe piRNA loss and widespread transposon derepression in the male germline, while leaving *Stellate* silencing intact (30, 31). Accordingly, *rhi* mutant males remain fertile, reproducibly siring dozens of progeny under laboratory conditions. In this regard, *rhi* mutations act as separation-of-function “alleles” of the piRNA pathway that are uniquely suited for studies of transposition, because they de-silence transposons in the male germline without de-silencing *Stellate*, thereby maintaining sufficient male fertility for offspring to be collected (30). We therefore used *rhi* mutant males to examine transposition upon piRNA loss.

### A mutation accumulation strategy for studying transposition after piRNA loss

Another challenge in studying transposon movement is that bona fide transposition events may be rare in a single generation. In *D. melanogaster*, the spontaneous transposition rate has been estimated at approximately 0.57 insertions per genome per generation (32). Although *rhi* mutations are expected to elevate transposition rates based on their strong effects on transposon expression, the magnitude of this effect on actual transposition is unknown. To address this challenge, we designed a mutation accumulation strategy analogous to the classic experiments of Terumi Mukai (33–35) and recent variants (36, 37), allowing transposition events to accumulate across generations. Importantly, this scheme propagated a single male fly each generation rather than a population, thereby relaxing selection that might otherwise efficiently purge deleterious transposon insertions. In every generation, the focal male was outcrossed to females from a separately maintained inbred stock, and dominant visible markers allowed each major chromosome to be tracked unambiguously (SI Appendix, Fig. S1). Because male *Drosophila* lack meiotic recombination, focal chromosomes could be readily distinguished from chromosomes inherited from the separately maintained inbred stock, and they remain genetically separate across generations. This enabled us to follow a full complement of nonrecombining haploid genome across generations via a single male germline lineage, including the Y chromosome and both major autosomes (Fig. 1D). Moreover, because the focal chromosomes were maintained strictly heterozygous, recessive deleterious transposon insertions were largely shielded from selection. This offers an advantage over mutation accumulation experiments based on full-sib matings, in which recessive deleterious alleles can be exposed to selection through homozygosity. Together, this design enabled the accumulation of transposition events on focal chromosomes across generations with minimal loss due to recombination or selection.

A central feature of our approach is incorporating *rhi* mutations into a Mukai-style mutation accumulation scheme (Fig. 1D). One of the two focal autosomes tracked through the male germline lineage across generations carried a *rhi* mutant allele, whereas the separately maintained inbred stock that supplied females for each generation was heterozygous for *rhi*. Consequently, the genotype of each focal male depended on which maternal *rhi* allele it inherited. Because the *rhi* mutation is recessive, this design enabled us to establish two male germline lineages from a single ancestral male, one with and one without functional Rhi-dependent piRNA biogenesis. In each subsequent generation, focal males were individually crossed to females from the same *rhi* heterozygous stock, allowing either *rhi* control or mutant males to be selected to continue the mutation accumulation experiment. Our experimental design therefore allowed transposition events to accumulate across generations in single male germline lineages derived from a common ancestry, either with or without functional *rhi* and piRNA-mediated transposon silencing (Fig. 1D).

To identify genome-wide transposon insertions, we adopted Nanopore long-read sequencing (38) from single flies (39, 40). We performed sequencing on females for two reasons. First, a single female yields more genomic DNA than a single male, as female flies are larger. Second, the highly repetitive nature of the *D. melanogaster* Y chromosome makes it recalcitrant to transposon insertion mapping, so we decided to focus on autosomal insertions in this work. Sequencing females avoids allocating reads to the large, repeat-rich Y chromosome, thereby increasing the proportion of informative reads available for detecting autosomal insertions. Using marked chromosomes, we isolated single females carrying focal autosomes from either the control or mutant lineage for Nanopore sequencing. Importantly, these females differed only in the two focal autosomes, which independently accumulated transposon insertions during the experiment, while all other chromosomes were effectively identical because they originated from the same separately maintained inbred stock (Fig. 1D).

If the piRNA pathway suppresses transposition, we would expect chromosomes inherited through the control male germline lineage to accumulate few, if any, new transposon insertions, whereas those inherited through the *rhi* mutant male germline lineage should accumulate substantially more insertions.

### Loss of *rhi* unleashes transposition in the male germline

We sequenced one fly from the *rhi* control lineage and one from the mutant lineage after 45 generations of divergence in *rhi* status, an experiment spanning over 2 years (Fig. 2A). Single-fly Nanopore sequencing of the two samples yielded 10- and 12-fold genome coverage, respectively, and the read N50 was about 14 kb for both samples (SI Appendix, Table S1). Given that most transposons in the *D. melanogaster* genome are 5-10 kb in length, the observed read-length distribution should allow a substantial fraction of reads to span entire transposon insertions. In other words, single reads often contained the complete transposon insertion, target site duplication (a molecular footprint often generated during transposition) (41–43), and flanking genomic sequences sufficient to resolve the insertion site. Using the software TLDR (44), we identified non-reference transposon insertions in flies from two lineages that accumulated mutations with or without *rhi* for 45 generations. These include non-reference insertions shared between lineages, which were inferred to be ancestral (Fig. 2B), as well as non-reference insertions unique to one lineage, some of which may represent new insertions acquired during the experiment (see below).

**Fig. 2.**
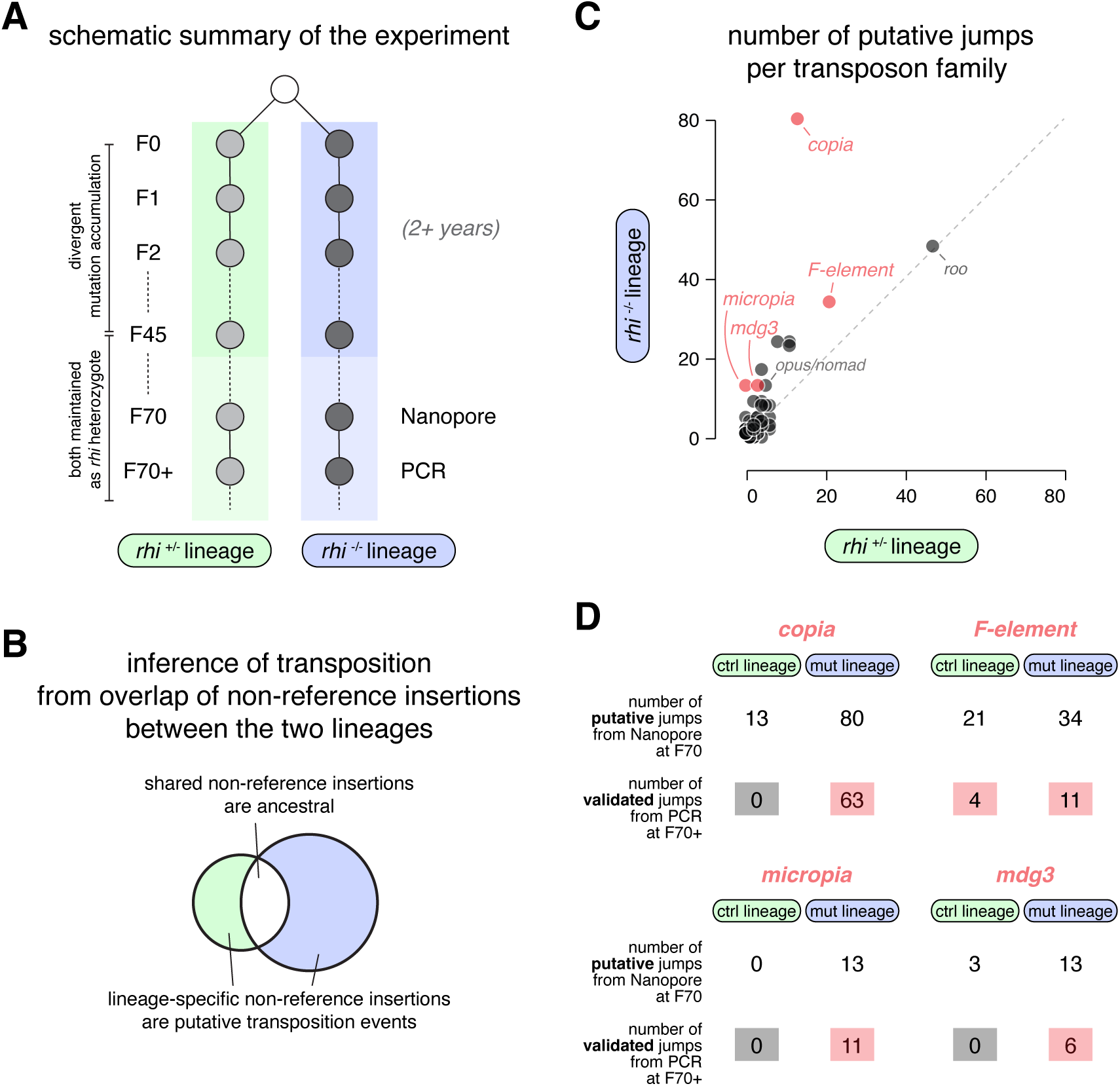
Loss of *rhi* unleashes transposition in the male germline. (A) A schematic summary of the experiment. (B) Inference of transposition from overlap of non-reference insertions between lineages. Shared non-reference insertions are ancestral, whereas lineage-specific non-reference insertions are putative transposition events. (C) Comparison of the number of putative jumps per transposon family in *rhi* control and mutant lineages. (D) Number of validated jumps from PCR for *copia, F-element, micropia,* and *mdg3*.

Grouping these lineage-specific non-reference insertions by transposon family allowed us to quantify putative transposition events for each family in the two lineages (Fig. 2C). We observed a similar or higher number of putative jumps in the *rhi* mutant lineage for most transposon families, among which one family, *copia*, appeared to transpose substantially more frequently in the *rhi* mutant male lineage, with 80 putative insertions in the mutant compared to 13 in the control. However, these “lineage-specific” insertions may include not only de novo insertions (true positives), but also ancestral insertions that were detected in one lineage but missed in the other for reasons such as limited sequencing depth (false positives). To independently validate these putative jumps, we used PCR to detect the junction sites of individual transposon insertions using genomic DNA extracted from flies in subsequent generations that inherited the mutation-accumulated focal autosomes. If an insertion produced a band in both lineages, it was classified as ancestral (false positive); conversely, if a band was detected in only one lineage, the insertion likely represented a genuine de novo event in that lineage (true positive). Applying this approach to all putative *copia* insertions, we validated 63 transposition events in the *rhi* mutant male lineage (Fig. 2D; SI Appendix, Table S3), indicating that *copia* produced at least 63 de novo insertions on two focal autosomes.

Because the two focal autosomes we tracked accounted for approximately one-third of the genome, we estimate that this approach recovered only a fraction of all transposition events. Strikingly, none of the putative *copia* insertions in the control lineage were validated (they were all false positives), suggesting that *copia* did not transpose at all in flies where the piRNA pathway was intact.

Why was *copia* so active transpositionally in the male germline? Testis RNA-seq showed that, in *rhi* mutants, *copia* produces the most abundant transposon mRNA (30)—an order of magnitude more than the next most highly expressed transposon family (Fig. 3A). Thus, the exceptional abundance of *copia* transcripts likely contributes, at least in part, to *copia* having the highest transpositional activity in the male germline.

**Fig. 3.**
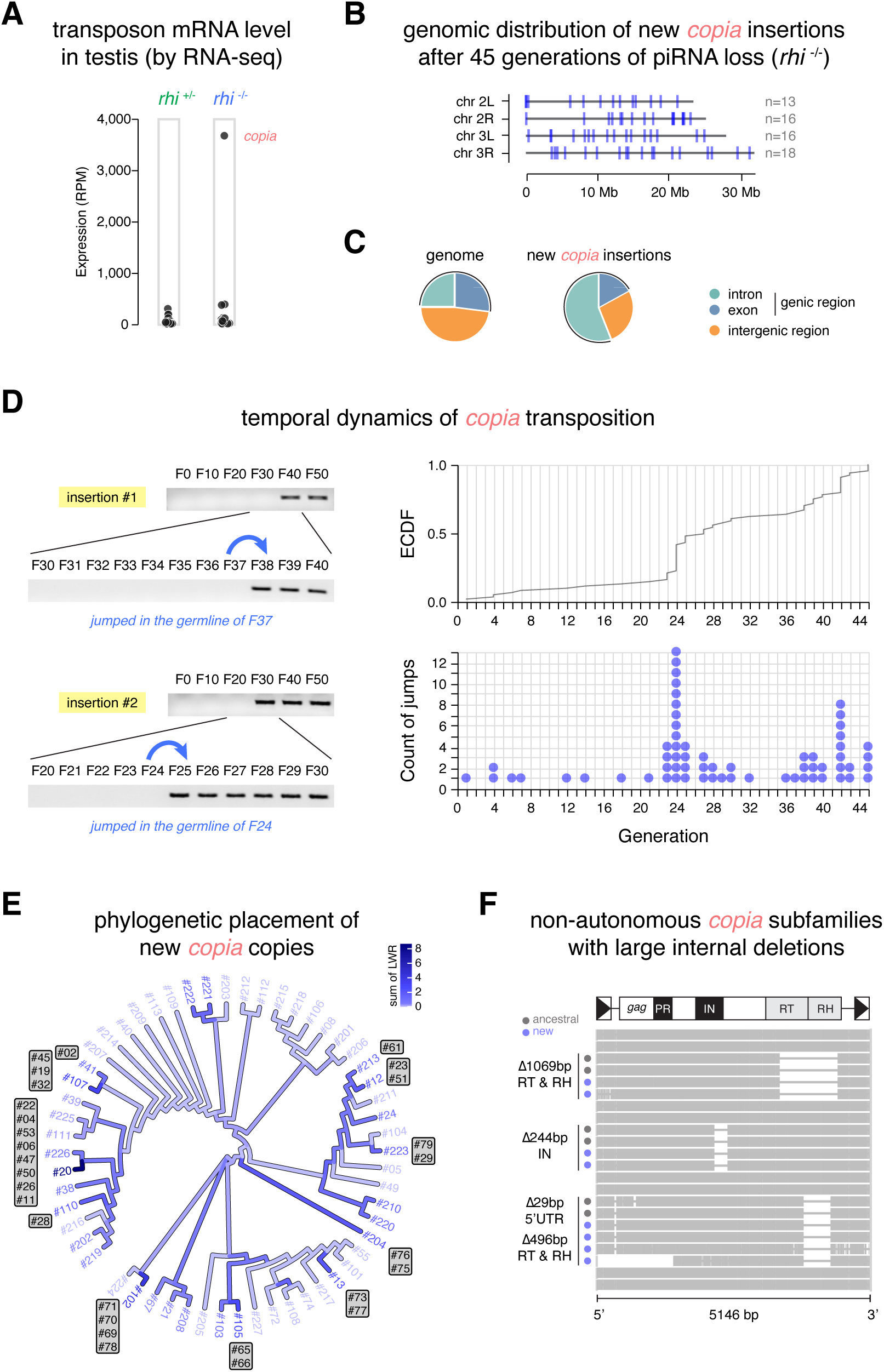
Tempo and mode of *copia* transposition. (A) Transposon mRNA levels in *rhi* control and mutant testes, showing the exceptional abundance of *copia* transcripts in *rhi* mutants. (B) Genomic distribution of new *copia* insertions after 45 generations of piRNA loss. (C) Makeup of the genome and distribution of new *copia* insertions with respect to genes. (D) Temporal dynamics of *copia* transposition. Left: example gel results first checking in which 10-generation range a given *copia* copy jumped and then checking at which exact generation the transposition occurred. Right: ECDF (empirical cumulative distribution function) and histogram of the number of *copia* jumps per generation, depicting bursts of transposition. (E) Phylogenetic placement of new *copia* copies. New insertions are copies detected by Nanopore sequencing in only one lineage and subsequently validated by PCR. The reference tree (blue) includes all the ancestral copies: those detected by Nanopore in both lineages (numbered 2xx), and those detected in only one lineage—control (numbered 1xx) or mutant (numbered <100)—but later found to be present in both lineages by PCR and thus reclassified as ancestral. Each new copy was initially provisionally placed on every branch of the reference tree, and the respective likelihood was computed to select a subset of promising branches. The precise maximum likelihood placements were then computed for the selected branches. Branches of the reference tree are colored by the sum of normalized placement likelihood across all new insertions. Branches with high summed likelihood likely gave rise to multiple descendant insertions, while branches with no summed likelihood may represent ancestral copies with no descendants. When a new insertion’s highest single-branch likelihood exceeded 50%, it is labeled with a two-digit number in a grey box adjacent to its most probable source copy. See also SI Appendix, Fig. S2. (F) Alignment of full-length *copia* and nonautonomous subfamilies with large internal deletions. Ancestral copies are marked with a gray circle, while new copies are marked with a blue circle.

We also examined putative transposition events from four additional transposon families: *micropia*, *F-element, mdg3, and opus*/*nomad* (Fig. 2D). After 45 generations of piRNA loss, we validated that *micropia* transposed 11 times, *F-element* transposed 11 times, *mdg3* transposed 6 times, and *nomad* showed no validated transposition events on the focal autosomes. Notably, we did not validate a single transposition event in the control lineage for most of these families, except for *F-element* which jumped 4 times. Collectively, these results provide direct evidence that the piRNA pathway suppresses male germline transposition to very low levels in the control lineage over 45 generations.

### Tempo and mode of *copia* transposition

Next, we characterized the genomic distribution of the new *copia* insertions. All 63 new *copia* insertions resided on autosomes (i.e., chromosomes 2 and 3) rather than sex chromosomes, independently validating our crossing scheme designed to accumulate mutations on autosomes. New *copia* insertions were distributed across chromosome arms on both focal autosomes, with the insertion numbers roughly scaling with chromosome-arm length (Fig. 3B). Relative to a random genomic distribution, *copia* insertions were enriched in genic regions (73% of insertions observed in genes vs. 52% expected under a random model (45); p < 0.001) (Fig. 3C). Within genes, *copia* insertions were enriched in introns relative to exons (3.2-fold enrichment, p < 0.001). Unless the integration machinery of *copia* can distinguish introns from exons, this pattern most likely reflects purifying selection against exonic insertions. Although selection was relaxed in our mutation-accumulation experiment, dominant deleterious exonic insertions would still have been subject to selection. Together, insertional preference for genic regions and selection against exonic insertions likely produced the overall enrichment of *copia* insertions in introns.

Our longitudinal experiment spanning 45 generations provided a unique opportunity to examine the temporal dynamics of *copia* transposition. Using genomic DNA collected at each generation, we dated all 63 *copia* insertions by performing PCR with junction-spanning primers (Fig. 3D). Once detected, each insertion persisted through all subsequent generations, consistent with our experimental design and supporting the interpretation of these events as bona fide transpositions on focal autosomes. During the first 22 generations, *copia* transposed at a low but relatively steady rate of approximately one insertion every 2-3 generations. Remarkably, a burst of transposition occurred around generation 24, with 13 insertions arising in a single generation. Subsequently, the transposition rate declined over several generations before a second, smaller burst at generation 42 (Fig. 3D). Overall, *copia* insertion counts per generation showed strong overdispersion relative to a constant-rate Poisson expectation, with variance exceeding the mean by more than threefold. This level of variability is readily apparent in the distribution across generations and is not consistent with a homogeneous transposition process. (Fig. 3D). We therefore conclude that, in the absence of piRNA control, *copia* transposes in temporally clustered bursts.

Given that full-length transposon sequences were available from long-read sequencing, we reconstructed the genealogy of *copia* insertions (Fig. 3E and SI Appendix, Fig. S2). We first generated a reference phylogeny of ancestral *copia* copies present before the start of mutation accumulation. Using a phylogenetic placement framework, we then inferred the most likely placement of each new *copia* insertion on the reference tree using EPA-ng (46). From this analysis, we found that transposition activity was distributed across multiple ancestral *copia* copies rather than concentrated within one or a few copies. To further investigate the origins of new insertions, we annotated *copia* copies whose maximum placement likelihood exceeded 50% next to their most probable parental copy on the tree. This analysis suggested that more than 10 ancestral *copia* copies gave rise to new progeny, with the most active copy serving as the probable source of 8 new insertions (Fig. 3E). In addition, the burst of 13 transposition events at generation 24 can be attributed to at least 6 source copies, rather than a single hot copy. Together, these results demonstrate that multiple *copia* copies are capable of transposition, echoing earlier work on *copia* (47), and arguing against a master-copy model in which transposition is dominated by one or a few highly active progenitor elements (48–50).

Although recent improvements have substantially increased Nanopore read accuracy, residual basecalling errors can still sometimes limit confidence in subtle phylogenetic differences among closely related sequences, especially when coverage is not sufficiently deep. We therefore asked whether large insertions or deletions could provide additional insight into *copia* transposition. Alignment of all *copia* insertions identified three subfamilies carrying distinct internal deletions (Fig. 3F): one with a 1069-bp deletion spanning the reverse transcriptase (RT) and RNase H (RH) domains, one with a 244-bp deletion in the integrase (IN) domain, and one with a 29-bp deletion in the 5′ UTR as well as a 496-bp deletion spanning RT and RH. The genealogy inferred from these deletions was fully consistent with the substitution-based phylogenetic placements described above, providing independent support for our conclusions. Importantly, these internally deleted subfamilies are nonautonomous because they cannot encode the full complement of proteins required to complete the *copia* replication cycle. Their recurrent transposition during the experiment suggests that nonautonomous *copia* copies can exploit RT, RH, and/or IN proteins supplied in trans by autonomous full-length copies. Thus, both autonomous and nonautonomous *copia* elements can transpose, consistent with prior reports of nonautonomous retrotransposon mobilization in mice and *Arabidopsis* (51–54).

In sum, we characterized the tempo and mode of *copia* transposition and found that multigenerational piRNA loss unleashes episodic bursts of transposition.

### Biological replicates uncover recurrent and stochastic features of transposition

The episodic transposition bursts observed in the original lineage raised a broader question of whether such dynamics is an intrinsic consequence of piRNA loss, or an idiosyncratic feature of a single fly lineage. To test this, we repeated the experiment with two independent biological replicates (A and B) from a distinct genetic background, initiated four years after the original experiment (Fig. 4A). Both replicates were derived from a single ancestral male and carried identical focal autosomes. We then accumulated transposon insertions on focal autosomes for 10 generations in each lineage, allowing direct comparison of transposition dynamics across independent evolutionary trajectories.

**Fig. 4.**
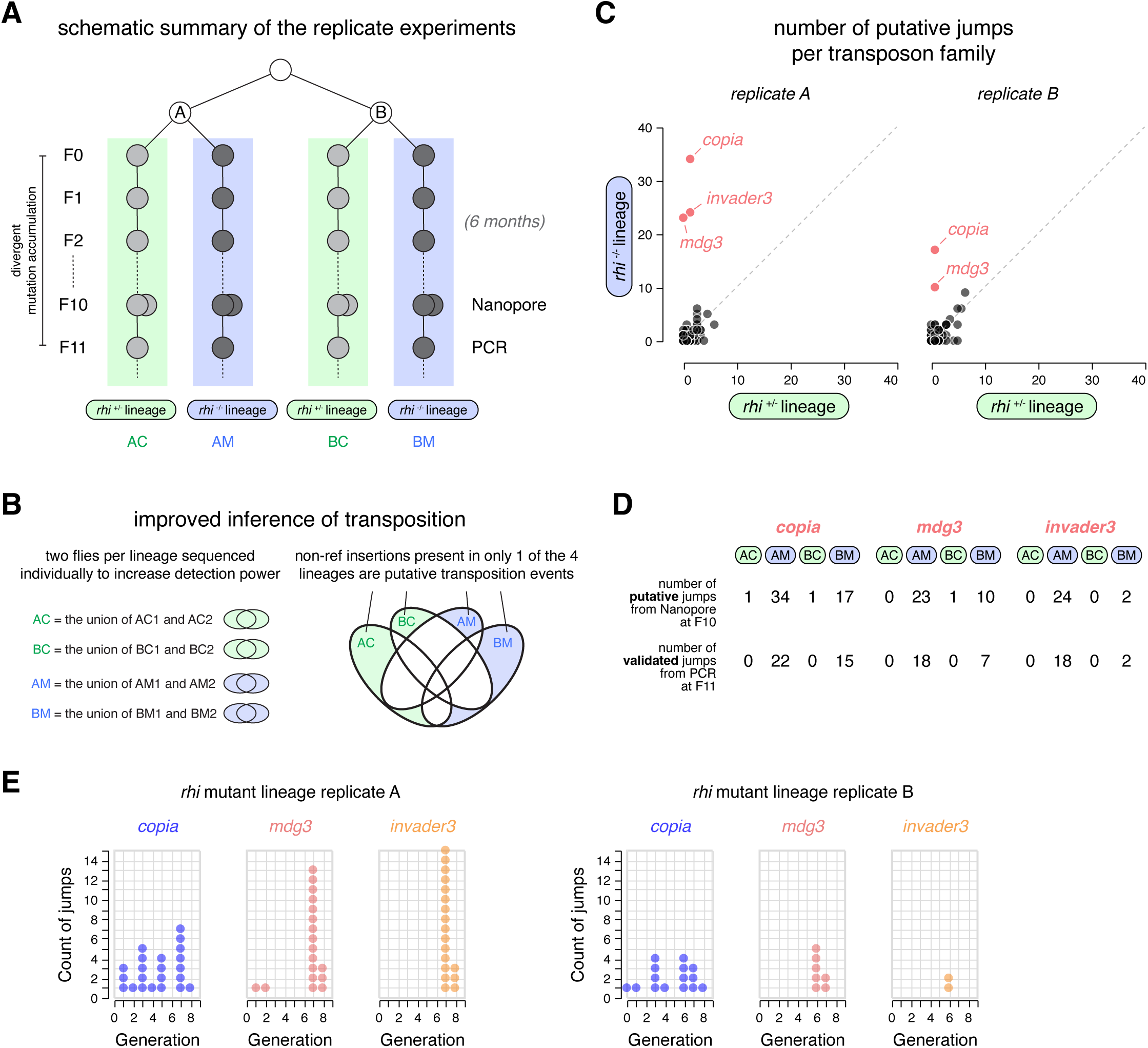
Biological replicates uncover recurrent and stochastic features of transposition. (A) A schematic summary of the replicate experiments. (B) Improved inference of transposition. Left: the union of insertions identified in two flies of a given lineage is used to increase detection power. Right: non-reference insertions present in only 1 of the 4 lineages are putative transposition events. (C) Comparison of the number of putative jumps per transposon family in *rhi* control and mutant lineages of both replicates. (D) Number of validated jumps from PCR for *copia, mdg3,* and *invader3*. (E) Histogram of the number of jumps per generation for *copia*, *mdg3,* and *invader3*.

To improve the inference of transposition events, we introduced two modifications in the replicate design (Fig. 4B). First, to increase detection power, we sequenced two flies per lineage, effectively doubling coverage relative to the original experiment (SI Appendix, Table S2). We defined the union of insertions detected in two flies as the set of insertions present in a given lineage. Second, we classified insertions shared across multiple lineages as ancestral, and those unique to a single lineage as candidate transposition events. For example, a candidate insertion in replicate A *rhi* mutants was required to be absent from the A control as well as both mutant and control lineages in replicate B. This framework is expected to reduce both false positives and false negatives in the detection of transposition events.

Strikingly, quantification of putative new jumps per transposon family in both replicates revealed a clear pattern even before PCR validation (Fig. 4C): transposition was largely confined to *copia*, *mdg3*, and *invader3*, which appeared to have jumped more after 10 generations of *rhi* mutation than in controls. Using PCR, we independently validated most (∼93%) of the *copia*, *mdg3*, and *invader3* insertions in both replicate mutant lineages (Fig. 4D; SI Appendix, Table S4), confirming reproducible activation of these families upon piRNA loss. These three families are among the most derepressed transposons in *rhi* mutant testes (*mdg3*: 36-fold, *invader3*: 18-fold, *copia*: 13-fold) (30), although some other derepressed transposons (e.g., *accord*: 17-fold) showed no evidence of transposition, suggesting that elevated mRNA abundance can, in some cases, lead to transposition. Among the transpositionally active families, the magnitude of activity differed between replicates, with more transposition events in replicate A than in replicate B across all three families, despite replicate B being sequenced approximately 1.45-fold more deeply. While *copia* and *mdg3* were previously known to be transpositionally active (55, 56), *invader3* was originally identified from repeat annotations of the fly genome (57, 58) but to our knowledge has not been previously shown to transpose. Our work therefore provides the first evidence that *invader3* is an element currently capable of transposition.

Once again, we did not observe a single transposition event for *copia, mdg3,* or *invader3* in the control lineage of either replicate. This lends further support to the idea that the piRNA pathway suppresses male germline transposition to a very low level.

Leveraging the transposition events of *copia*, *mdg3*, and *invader3* in the replicate lineages, we asked whether episodic transposition is a generalizable consequence of piRNA loss. Using PCR, we dated every *copia, mdg3,* and *invader3* insertion in two replicate lineages (Fig. 4E). As observed previously for *copia* in the first experiment, all three families transposed in episodic bursts rather than at constant rates across generations. Episodic dynamics were especially pronounced for *mdg3* and *invader3*, which completed most mobilization within narrow temporal windows: in replicate A, 89% of new *mdg3* insertions and all of new *invader3* insertions occurred at generations 7 and 8; in replicate B, all of new *mdg3* and *invader3* occurred at generations 6 and 7. These observations suggest that: within individual replicate lineages, *mdg3* and *invader3* bursts were strongly correlated temporally, yet burst timing differed slightly between the two replicate lineages (Fig. 4E). The correlated behavior of *mdg3* and *invader3* may reflect their close evolutionary relationship (57), as related elements may undergo coordinated mobilization. Alternatively, this may reflect temporal windows of particular vulnerability in host control that permit transposition. Altogether, our findings support a model in which piRNA loss creates a permissive state for transposon mobilization, but the onset, magnitude, and duration of individual bursts emerge stochastically across lineages.

## DISCUSSION

The current paradigm in the field posits that the piRNA pathway represses transposon expression to protect the genome from transposition. Yet, most prior work has used transposon expression as a proxy for transpositional activity, leaving the actual impact of the piRNA pathway on transposon movement largely unaddressed. In this work, we developed a genetically tractable system that enabled direct longitudinal analysis of transposition with or without an intact piRNA pathway, while minimizing recombination and selection. Across three biological replicates, we found that *copia* repeatedly exhibited the highest transpositional activity in the *Drosophila* male germline, highlighting its strong mobilization when the piRNA pathway is disrupted (59). In contrast, we validated very few transposition events in the control lineage, firmly supporting the idea that the piRNA pathway tightly suppresses germline transposition.

Of note, we did not identify transposition events of families known to transpose specifically into highly repetitive regions of the genome, including R1/R2 in rDNA arrays (60) and HeT-A/TART/TAHRE in telomeres (61, 62). This could merely reflect our inability to confidently call insertions in highly repetitive genomic regions or a genuinely low transpositional activity of these elements in our experimental system, a distinction that will require future work. It is also important to note that, while our experimental scheme relaxes selection, dominant deleterious insertions are expected to be underrepresented, so there is still some degree of ascertainment bias in our detection of transposition events.

While previous studies have examined transposition following piRNA pathway disruption using other experimental systems or approaches (6–13), these methods have a limited ability to distinguish ancestral from new insertions, resolve lineage-specific dynamics, or capture rare or temporally clustered transposition events. As a result, transposition rates were often estimated by treating non-reference insertions as de novo events, or averaged across heterogeneous germline populations, potentially obscuring substantial variation in activity across individual lineages and time. In this work, we addressed these limitations using a mutation accumulation framework that enables longitudinal tracking of endogenous transposons on defined chromosomes within single germline lineages. This design allows direct inference of transposition events and fine-scale reconstruction of their temporal dynamics across generations. These features were essential for revealing that transposition following piRNA pathway disruption occurs in episodic, lineage-specific bursts, which are difficult to detect in pooled or short-term experimental designs.

Our work also provides additional insights into transposon movement. We demonstrated the transpositional activity of several families in the male germline, including *copia, micropia*, *mdg3,* and notably, *invader3*, whose transpositional activity had not been demonstrated previously. Also, full-length sequences of transposon insertions from Nanopore long-read sequencing allowed us to reconstruct the genealogy of individual transposon families and infer the transpositional activity of nonautonomous subfamilies that bear large internal deletions. We found that many ancestral copies of *copia* are transpositionally active, indicating that *copia* mobilization is distributed across multiple active copies (47) rather than dominated by a few master elements, as proposed for LINE-1 and Alu in mammals (48–50).

Perhaps most intriguingly, we demonstrated punctuated episodes of transposition after collapse of the piRNA pathway. These episodic bursts of transposon movement over time suggest that elevated transposon expression alone is not sufficient to predict the timing or magnitude of transposition. Notably, we observed a delay in the onset of transposition bursts across all three replicates, indicating that additional factors likely govern the initiation of transposition bursts. The transposition rate also often fell after a transient peak, indicating that it does not simply scale with the total transposon copy numbers in the genome, at least on the timescale of our experiments.

Disentangling intrinsic stochasticity of transposon activity from alternative explanations will require sampling more independent lineages, ideally including multiple siblings at every generation. If burst timing correlates across most or all independent lineages, environmental variation is the likely driver. If it correlates among siblings from a single parent but not across independent lineages, lineage-specific molecular changes are implicated. If no temporal correlation exists at either level, variation across single germ cells—individual sperm or oocytes—is the most likely explanation. Although technically challenging at single-cell resolution, monitoring mRNAs, small RNAs, and epigenetic marks throughout the course of the experiments could help resolve the molecular basis for the delay, initiation, and cessation of transposition bursts.

Importantly, if transposons mobilize in bursts, substantial variation can appear over very short timescales (63), fueling evolution with a large amount of mutational input in a temporally heterogeneous manner. In fact, such saltatory character of transposition has been described in *Drosophila* for *copia* (64–67) and *P-element* (68) in the 80s, but the exact cause was not known. Our work shows that perturbation of the piRNA pathway may have enabled these transposition bursts. Similar bursty dynamics have also been reported in plants (67, 69–71) and fungi (72, 73), suggesting that episodic transposition is a widespread feature across taxa. By dramatically reshaping the timing and magnitude of mutational input, bursts of transposon mobilization (triggered by genetic or environmental perturbations to transposon control mechanisms) could provide one molecular basis for contingency (74) and punctuation (75) in evolution.

## ACKNOWLEDGEMENTS

We thank Dan Barbash, Kevin Wei, Chris Ellison, Katelyn Boese, Ben McCormick, and members of the Clark Laboratory and the Feschotte Laboratory for discussions. We appreciate Bernard Kim and Hannah Gellert for advice on extraction of high molecular weight DNA from single flies, and Satyam Srivastav for advice on calling transposon insertions from Nanopore data. We are grateful to Grace Yuh Chwen Lee, Mariana Wolfner, and Daniel Siqueira de Oliveira for comments on the manuscript draft. P.C. is supported by the HHMI Hanna H. Gray Fellowship. This work was supported by grants from the National Institutes of Health (R01 GM097363 to A.A.A., R01 HD059060 to A.G.C. and M. Wolfner, and R35 GM122550 to C.F.).

## MATERIALS AND METHODS

### Fly stocks and crosses

Double balanced *rhi* mutant alleles—*w* [-]; *rhi* [2] / *CyO*; *Ki* / *TM3, Sb, Ser* and *w* [-]; *rhi* [KG], *w* [+] / *CyO*; *Ki* / *TM3, Sb, Ser*—were used in our previous work (30) and were originally gifts of Bill Theurkauf. The two *rhi* alleles could be distinguished because one of them is linked with *w* [+], which alters the eye color in the *w* [-] background. The *rhi* [2] stock was maintained as a separate inbred stock throughout the experiment, and it is always heterozygous because the chromosome carrying *rhi* [2] has acquired a recessive lethal background mutation.

Two focal autosomes were used for mutation accumulation, one of which was the chromosome 2 that carried *rhi* [KG]. To introduce an unmarked chromosome 3 as the other focal autosome for mutation accumulation, we crossed the double balanced *rhi* [KG] stock to a *w* [-] stock with no autosomal markers, generating a single male of the genotype w [-] / Y; *rhi* [KG], *w* [+] / +; + / *TM3, Sb, Ser*. This male is shown as the initial male in Fig. 1 and SI Appendix, Fig. S1. Crossing this male to four females from the double balanced *rhi* [2] stock, we generated two F0 males, one being a *rhi* heterozygote (control), *rhi* [KG] / *CyO*; + */ TM3, Sb, Ser*, and the other being a *rhi* trans-heterozygous mutant, *rhi* [KG] / *rhi* [2]; + */ TM3, Sb, Ser*. These two males initiated two lineages, one with a functional *rhi* allele and the other without. By crossing these two males individually to four females from the *rhi* [2] stock, males of the same genotype could be obtained at every generation.

Importantly, this cross scheme enforced a unique inheritance pattern of chromosomes. A (nearly) full complement of the nonrecombining haploid genome was inherited together: a chromosome Y, the chromosome 2 bearing *rhi* [KG], and an unmarked chromosome 3 were inherited together exclusively through the focal male at every single generation. Meanwhile, a chromosome X, a chromosome 2 that either carries *rhi* [2] or *rhi* [+] (i.e., *CyO*), and a chromosome 3 marked by *Sb* and *Ser* (i.e., *TM3*) were introduced from the mother, stayed in the focal male for one generation, and were then replaced by another copy of the same set of chromosomes from the *rhi* [2] stock. Thus, the focal autosomes were tracked across generations and accumulated mutations with or without *rhi*. We did not track the tiny chromosome 4 in this work, which accounts for <1% of the genome.

Because males are subfertile without *rhi*, we established up to three identical crosses (when possible) at each generation for both lineages. The cross that produced three males of the desired genotype was randomly selected to continue the lineage, whereas the other two crosses were discarded. This approach ensured that males of the desired genotype were obtained at every generation to continue the experiment.

For the replicate experiments, we used a different chromosome 3 from a different *w* [-] stock to accumulate mutations. The focal chromosome 2 carrying *rhi* [KG] was in principle the same as the original experiment, but it had likely acquired mutations after four years of independent passage in the lab. For the original experiment, after 45 generations of divergent mutation accumulation, both lineages were kept as *rhi* heterozygotes for convenience until generation 70, when the fly was sequenced. For replicate experiments, we sequenced F10 after 10 generations of divergent mutation accumulation.

### Nanopore long-read sequencing from single flies

We followed a recently described method (39, 40) to extract high molecular weight DNA from single flies for Nanopore sequencing. Briefly, the fly was homogenized with a pestle and then subject to proteinase K and RNase A digestion at 50 °C for 3 h. Next, we performed phenol-chloroform extraction of genomic DNA from the lysate. Using Dow Corning High Vacuum Grease, the aqueous and organic phases were separated by the grease, and the upper aqueous phase was transferred by pouring without pipetting. Genomic DNA was precipitated in ice-cold ethanol, producing a pellet that was washed in 80% ethanol, air dried, and resuspended in 30 µL of 10 mM Tris-HCl (pH 8.0) at 50 °C for 1 h. The solution was kept at 4 °C for a week to obtain proper resuspension before sequencing. Nanopore long-read sequencing was done by Plasmidsaurus. Summary statistics of the Nanopore data can be found at SI Appendix, Tables S1 and S2.

### Calling non-reference transposon insertions from Nanopore data

We mapped Nanopore reads to the dm6 (76) reference genome using minimap2 (77) - ax map-ont. Next, we used the software TLDR (44) to identify non-reference transposon insertions, with the reference genome dm6 (76) and a transposon consensus library downloaded from GitHub (https://github.com/bergmanlab/drosophila-transposons). We focused on the insertion calls passing TLDR’s quality checks and having median MapQ >20, although results were similar without these filters. All calls were supported by at least one read spanning the entire insertion (78). Target site duplications and exact insertion sites inferred by TLDR were not always accurate and therefore were manually curated.

### Identification and validation of de novo transposition events

For the original experiment, we designated non-reference transposon insertions present in only one of the two lineages as putative de novo transposition events. For the replicate experiments, we required that a non-reference transposon insertion be present in one and only one of the lineages to be putative de novo transposition events. To validate the putative jumps, we designed primers targeting the junction of each insertion, with one primer within the transposon sequence and the other in the flanking genomic region. PCR products ranged from 500-1000 bp. For the original experiment, we used genomic DNA from generations 73-75 for PCR. For the replicate experiments, we used genomic DNA from generation 11 for PCR. For a given insertion, if a product was amplified in both control and mutant lineages, the insertion was classified as ancestral; if amplified only in one lineage, it was classified as a de novo insertion in that lineage; if no product was amplified in either lineage, the insertion was excluded from the analysis. Summary of the validation results can be found at SI Appendix, Tables S3 and S4. False-positive transposition events often arise from failure to detect ancestral insertions in one of the lineages rather than from spurious insertion calls. In the original experiment, the two samples each failed to detect 7 and 9 ancestral *copia* insertions, respectively (8 on average) (SI Appendix, Tables S3). In contrast, in the replicate experiments, the four lineages failed to detect 1-4 ancestral insertions across *copia*, *mdg3*, and *invader3* (2 on average) (SI Appendix, Tables S4), suggesting that the improved experimental design reduced the rate of false-positive transposition calls.

### Dating transposition events

For all validated transposition events, we sought to identify the generation in which they arose. For the original experiment with 45 generations of divergent mutation accumulation, we first performed PCR on genomic DNA from every 10 generations (F0, F10, F20, F30, F40, and F50). Once we identified the 10-generation interval in which a given insertion first appeared, we then did PCR with genomic DNA extracted at every generation within that interval. For the replicate experiments with 10 generations of divergent mutation accumulation, we directly performed PCR on genomic DNA from every generation. Overall, this approach confirmed that, once detected, each insertion was present in all subsequent generations, consistent with the expectation of bona fide transposition events in our experimental design.

### Analysis of transposon expression in fly testes

RNA-seq data of *rhi* mutant and control testes were published previously (30).

### Alignment of *copia* copies

We extracted full-length transposon sequences from the consensus assembled from Nanopore reads by TLDR and used mafft (--auto –adjustdirection) to generate a multiple sequence alignment (MSA). Visual inspection of the MSA identified several subfamilies with identical large internal deletions, suggesting that they are nonautonomous elements.

### Phylogenetic placement of *copia* copies

We reconstructed a MSA with the ancestral *copia* copy sequences using mafft and removed poorly aligned columns using trimAL (79) - gappyout. A reference phylogenetic tree was constructed using the trimmed ancestral sequence MSA with RAxML-ng v1.2.0, employing the GTR+G+F substitution model and 200 bootstrap replicates. The MSA of new insertions, together with the reference tree and the estimated substitution model parameters, was used as input for EPA-ng (46) v0.3.8 for phylogenetic placement. EPA-ng computes the maximum likelihood placement of each query sequence on candidate branches and outputs normalized likelihood (likelihood weight ratio, LWR) in a .jplace file. For each new copia insertion *i*, we denote the normalized placement likelihood on branch *j* as *p_ij_*, where *j* ∈ *L_i_* represents the set of candidate branches evaluated for sequence *i*. By construction, ∑*_j_ p_ij_* = 1. The sum of likelihood ratios for branch *j*, *PL_j_*, was calculated as *PL_j_* = ∑*_i_ p_ij_*, which is used to color the branches of Fig. 3E. All of the *copia* sequences and the codes for both phylogenetic placement and visualizations are available at GitHub (https://github.com/perryXuu/copia-phylogenetic-placements).

### Data visualization

Most data visualization was done in Python 3 using JupyterLab and the following software packages: numpy, pandas, and altair. Multiple sequence alignments were visualized using the NCBI MSA Viewer (https://www.ncbi.nlm.nih.gov/projects/msaviewer/). The phylogenetic tree was visualized using the ggtree (80) package in R.

### Data, Materials, and Software Availability

The sequencing data generated in this study are available at NCBI SRA under the accession PRJNA1521616 (81).

