## Supplementary material for "piRNA loss unleashes episodic transposition bursts": SI Appendix

**This PDF file includes:**

Figures S1 to S2

Tables S1 to S4

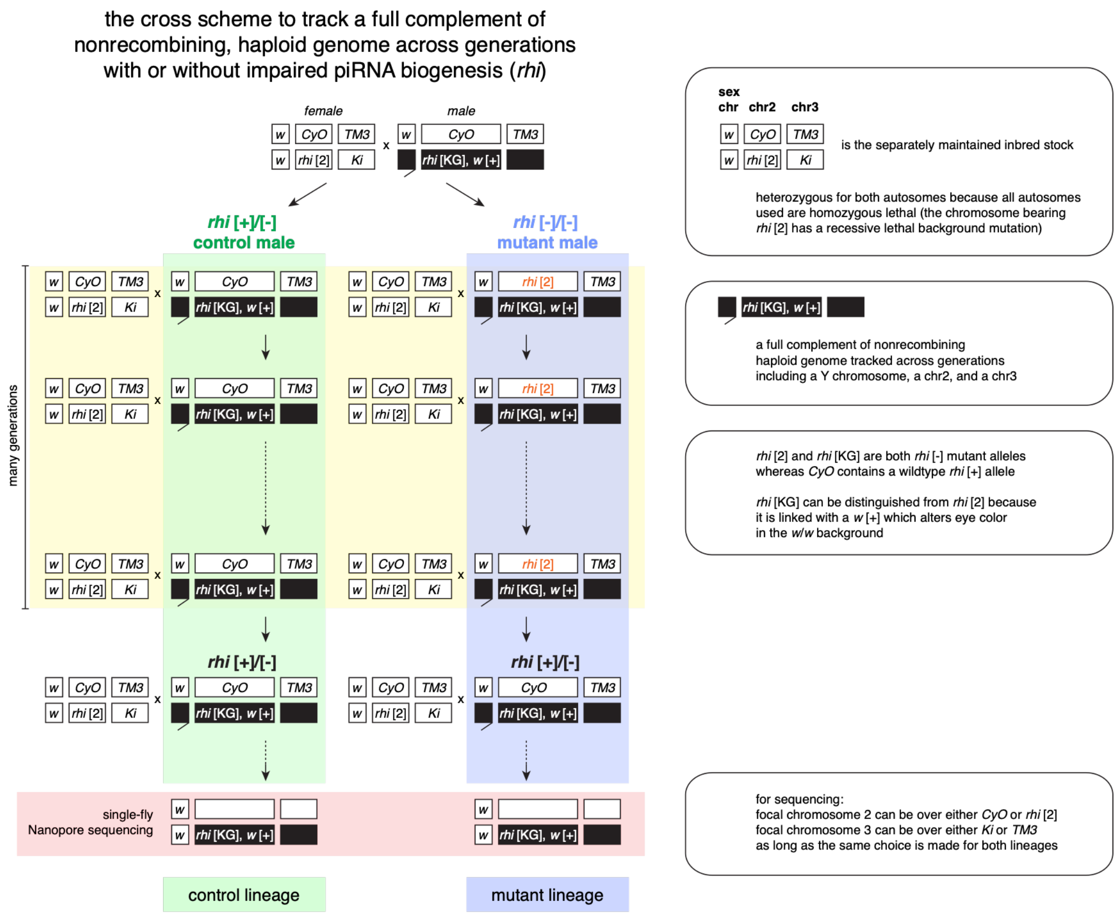

**Fig. S1. A mutation accumulation strategy for studying transposition with or without Rhi-dependent piRNA biogenesis.**

The same cross scheme as Fig. 1D with detailed genotype labels for almost every chromosome in the experiment. Additional explanatory texts are included on the right.

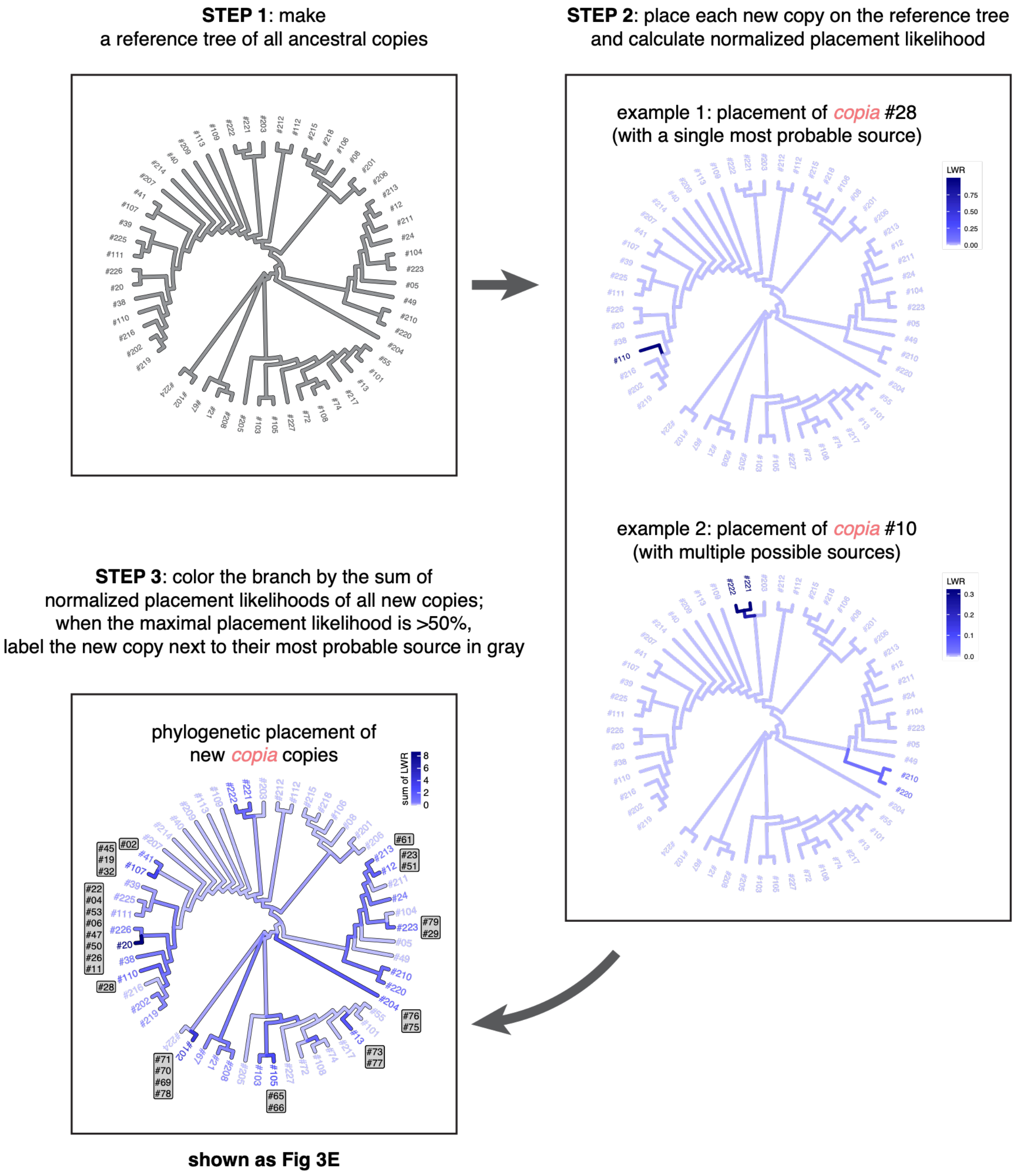

**Fig. S2. Procedures for phylogenetic placement of new copia copies.**

Step-by-step procedures to generate the plot shown in Fig. 3E.

**Table S1.** Summary statistics of the Nanopore data for the original experiment.

| Nanopore stats | control lineage | mutant lineage |
|---|---|---|
| **Total bp sequenced** | 1,297,810,198 | 1,548,690,601 |
| **Total number of reads** | 333,372 | 330,649 |
| **reads N50** | 13,892 | 14,021 |
| **Longest read** | 142,558 | 132,586 |
| **Estimated coverage** (assuming 130Mb genome) | 10x | 12x |

**Table S2.** Summary statistics of the Nanopore data for the replicate experiments.

| Nanopore stats | AC | | AM | | BC | | BM | |
|---|---|---|---|---|---|---|---|---|
| | AC1 | AC2 | AM1 | AM2 | BC1 | BC2 | BM1 | BM2 |
| **Total bp sequenced** | 3,242,312,613 | 2,132,318,890 | 1,927,131,951 | 1,544,960,275 | 2,987,684,036 | 4,129,188,232 | 1,713,103,495 | 3,328,441,877 |
| **Total number of reads** | 411,888 | 310,376 | 343,130 | 162,906 | 315,370 | 856,100 | 240,469 | 707,520 |
| **reads N50** | 17,355 | 15,529 | 13,469 | 23,990 | 21,579 | 8,377 | 14,691 | 9,205 |
| **Longest read** | 155,493 | 138,284 | 164,983 | 153,823 | 149,680 | 184,331 | 146,371 | 132,933 |
| **Estimated coverage** (assuming 130Mb genome) | 25x | 16x | 15x | 12x | 23x | 32x | 13x | 26x |

**Table S3.** Summary of the transposition validation results for the original experiment.

| | control lineage | | | | mutant lineage | | | |
|---|---|---|---|---|---|---|---|---|
| | copia | F-element | micropia | mdg3 | copia | F-element | micropia | mdg3 |
| **# putative jumps from Nanopore** | **13** | **21** | **0** | **3** | **80** | **34** | **13** | **13** |
| **# validated jumps from PCR (i.e., a band only in this lineage)** | **0** | **4** | **0** | **0** | **63** | **11** | **11** | **6** |
| **# ancestral insertion from PCR (i.e., a band in both lineages)** | 9 | 10 | 0 | 1 | 7 | 8 | 1 | 3 |
| **# inconclusive from PCR (i.e., no band)** | 4 | 7 | 0 | 2 | 10 | 15 | 1 | 4 |

**Table S4.** Summary of the transposition validation results for the replicate experiments.

| | AC | | | AM | | | BC | | | BM | | |
|---|---|---|---|---|---|---|---|---|---|---|---|---|
| | **copia** | **mdg3** | **invader3** | **copia** | **mdg3** | **invader3** | **copia** | **mdg3** | **invader3** | **copia** | **mdg3** | **invader3** |
| **# putative jumps from Nanopore** | **1** | **0** | **0** | **34** | **23** | **24** | **1** | **1** | **0** | **17** | **10** | **2** |
| **# validated jumps from PCR (i.e., a band only in this lineage)** | **0** | **0** | **0** | **22** | **18** | **18** | **0** | **0** | **0** | **15** | **7** | **2** |
| **# ancestral insertion from PCR (i.e., a band in both lineages)** | 1 | 0 | 0 | 4 | 0 | 0 | 0 | 1 | 0 | 0 | 2 | 0 |
| **# inconclusive from PCR (i.e., no band)** | 0 | 0 | 0 | 8 | 5 | 6 | 1 | 0 | 0 | 2 | 1 | 0 |
| | 22+18+18+15+7+2 = 82 validated transposition events in two mutant lineages<br>82 / (82+4+2) ≈ 93% validation rate in two mutant lineages (excluding the inconclusive ones) | | | | | | | | | | | |
